# Single cell variations in expression of codominant alleles A and B on RBC of AB blood group individuals

**DOI:** 10.1101/439083

**Authors:** Anjali Bajpai, Vidya Jonnalagadda, Badanapuram Sridevi, Pathma Mutthukotti, Rachel A Jesudasan

**Affiliations:** Center for Cellular and Molecular Biology, Habsiguda, Hyderabad –500 007, T.S. India; Bhavan’s Vivekanada College, Sainikpuri, Secunderabad – 500094. T.S

**Keywords:** ABO locus, codominant alleles, mosaicism, differential expression, immunofluorescence

## Abstract

One of the key questions in biology is whether all cells of a “cell type” have more or less the same phenotype, especially with relation to non-imprinted autosomal loci. Recent studies point to differential allelic expression of autosomal genes being a prevalent phenomenon responsible to confer phenotypic variability at individual cell level. However, most studies have been carried out in actively transcribing cells. Here we display cellular mosaicism arising from differential allelic expression for the cell surface glycoprotein in the enucleated RBCs. We studied the expression of the A and B histo-blood group antigens encoded by the co-dominant alleles in individual RBCs using immunofluorescence. We assessed the relative levels of the co-dominant alleles I^A^ and I^B^ in 2512 RBC from 24 individuals with AB blood group using Cy3- and FITC- tagged antibodies. Quantification of individual fluorescence intensities from each cell and test of their normal distribution revealed that contrary to the general belief that all RBC in AB individuals express both antigens in comparable amounts, they segregated into 4 groups: showing normal distribution for both antigens, either antigen, and neither antigen; the deviation from normal distribution could not be correlated to maternal/paternal origin, thus appear to be stochastic. Surprisingly, very few people showed any correlation between the amounts of these two antigens on RBC. In fact, the ratio of antigen A to B in the entire set of samples spanned over 5 orders of magnitude. This variability in amount of the antigens A and/or B, combined with a lack of correlation between the amounts of these two antigens resulted in unique staining patterns for RBC, generating widespread mosaicism in the RBC population of AB blood group individuals.

## Introduction

Fitness of an organism is determined by the phenotypic and cellular heterogeneity, which in turn arises from gene expression variations(Espinosa-Soto et al., 2011; Bódi et al., 2017). Whole-genome studies(Lo et al., 2003; Gimelbrant et al., 2007; Zwemer et al., 2012; Deng et al., 2014) have uncovered presence of remarkable phenotypic variations arising due to differential expression of alleles in individual cells. While imprinted genes represent extreme examples of differential allele activity, recent studies point to allelic variations being a more prevalent phenomenon responsible to confer striking phenotypic variability (see review (Eckersley-Maslin and Spector, 2014; Reinius and Sandberg, 2015; Chess, 2016; Gendrel et al., 2016)). The choice of the allele expressed could be random or could be fixed and stably inherited in mitotic clonal population (Reinius et al., 2016). The pervasiveness of random monoallelic expression has been observed across various cell types that display active transcription. Interestingly, random monoallelic expression was found to increase with cell differentiation during development(Eckersley-Maslin et al., 2014) and is enriched for tissue-specific genes including cell surface receptors (Deng et al., 2014; Branciamore et al., 2018). Here we have looked at expression of co-dominant alleles of the ABO gene locus (Yamamoto et al., 1990), that results in the formation of antigen A or antigen B, on surface of reticulocytes and most of the epithelia, barring some such as muscle cells and neurons (Clausen and Hakomori, 1989; Kominato et al., 2020).

Alleles I^A^ and I^B^ that are responsible for the production of antigens A and B(Tuppy and Staudenbauer, 1966; Ginsburg, 1972; Yamamoto et al., 1990; Watkins and Morgan, 1993) respectively, of the ABO blood group are cited as classic examples of co-dominant expression of Mendelian traits(Von Dungern and Hirschfeld, 1911). The A and B antigens are produced by the differential glycosylation of a common substrate, H, present on the surface of all human cells. Allele I^A^ encodes α1-3 N-acetylgalactosaminyl transferase which adds an N-acetyl galactosamine residue to H disaccharides present on glycolipids or glycoproteins; this modified form is termed antigen A. Allele I^B^ encodes α1-3 galactosyl transferase which adds a galactose residue at the same position of the H antigen; this modified form is termed antigen B. Null alleles at this locus are termed i, and the unmodified substance H is termed O antigen. Alleles I^A^ and I^B^ are both dominant over allele i, but are codominant with respect to each other. Recently, natural and engineered enzymes with dual specificity (cis-AB alleles) have been reported that can modify H antigen to produce both A and B antigens. However, an individual molecule of substance H can be modified to produce either A antigen or B antigen, but not both.

Cells of individuals with genotype I^A^I^B^ express both, A antigen and B antigen, on their surfaces. It is generally believed that these two antigens are present in more or less equivalent amounts, as judged by agglutination reactions. Certain types of I^A^ and I^B^ alleles are known to be ‘weak’, either due to lower level of transcription and/or poor catalytic activity of the encoded polypeptide product (Yamamoto et al., 1993), (Chester and Olsson, 2001). For example, I^A^ and I^B^ alleles containing missense mutations showed variation in expression when transfected into HeLa cells in terms of mean antigen amount as well as number of antigen expressing cells (Seltsam and Blasczyk, 2005). Nevertheless, one would expect that all the cells within an I^A^I^B^ individual would be fairly homogenous in terms of the stoichiometric ratios of the A and B antigens.

Here we report the quantification by immunofluorescence, of A and B antigens on 2512 peripheral red blood cells (RBC) from 24 individuals of AB blood type. We observed that a majority of the subjects (20 of the 24) in this cohort did not have RBC that stained equally for antigens A and/or B; in most cells, each RBC stained substantially more for either antigen A or antigen B. Quantification of both antigens on each cell revealed a wide variation in the amount of A and/or B antigen in most samples. Of the 24 individuals tested, only 8 showed appreciable or strong positive correlation (Carl Pearson correlation coefficient r > 0.5). The effect of these two levels of variation in quantity of surface antigens is that most individuals of AB blood group in this study group were phenotypic mosaics for these proteins. These observations challenge the presumption of stoichiometric expression of these two alleles within the RBC population and may be indicative of epigenetic mechanisms that can render these two alleles functionally non-equivalent within a cell.

## Materials and Methods

### Ethics Statement

Blood samples were collected with informed consent of the subjects, after obtaining clearance (IEC-51/2016) from the ethical committee, Centre for Cellular and Molecular Biology (CCMB), Hyderabad, India. The subjects were healthy donors from the institute. All the experimental procedures followed were as per strict guidelines and regulations issued by ethical committee, CCMB.

#### Immunofluorescent detection of antigens A and B on RBC

100 μl of peripheral blood was fixed on poly-lysine coated cover slips using 3.7% formaldehyde. The cells were blocked with 2% BSA and incubated serially in (a) mouse monoclonal Anti-A antibody (1:10 dilution; Tulip Diagnostics (P) Ltd. India) for 90 min., (b) Cy3-tagged anti-mouse antibody (1:500 dilution; Amersham Pharmacia) for 60 min, (c) mouse monoclonal Anti-B antibody (1:10 dilution; Tulip Diagnostics (P) Ltd. India) for 90 min, and (d) FITC tagged anti-mouse antibody (1:200 dilution; Vector Labs). After each incubation, the slides were washed in 1x PBS (Phosphate Buffered Saline) for 15 min. After the final wash, slides were mounted using a glycerol-based anti-fade reagent, Vectashield (Vector Labs). Control reactions carried out to rule out non-specific binding were incubation of (a) anti-A antibody to RBC of O and B blood group types, (b) anti-B antibody to RBC of O and A blood group types, and (b) antimouse secondary antibody to RBC of all four blood groups (O, A, B, and AB) in the absence of any primary antibody. All procedures were done at room temperature.

#### Image capture

Slides were scanned at 40x (dry) and 100x (oil) magnifications using the dual scanning, multi-track mode of a confocal microscope (Zeiss, LSM510 META). The numerical aperture used was 0.95 for 40x and 1.4 for 100x. Images containing 5 Z-stacks were captured for quantification (100x magnification) on Axioplan-2 using Axiovision version-4 camera (Carl Zeiss). The intensity of fluorescence of FITC and Cy3 was quantified from the projected images using the LSM-FCS Version 3.2SP software (Carl Zeiss) and corrected for background fluorescence. Typically, around 100 cells were analyzed from each sample slide.

#### Statistical analysis

Data were analyzed using MS Excel and R Statistical Software. Outliers in box plots were identified using default parameters of R. The Shapiro Wilk test was used to determine the probability that a given set of fluorescence values conformed to a normal distribution. For each subject, the Karl Pearson correlation coefficient was calculated using basic functions in Excel.

### Data availability statement

The authors affirm that all data necessary for confirming the conclusions of the article are present within the article, figures, and tables.

## Results and discussion

### Unequal staining for A and B antigens on RBC from AB blood group individuals

The RBC from 24 (15 males and 9 females) AB blood group individuals were fixed on slides and incubated with anti-A antibody detected by Cy3 (red fluorescence) tagged secondary antibodies and then incubated with anti-B antibody detected by FITC (green fluorescence) tagged secondary for antigen B. Contrary to expectations of finding equivalent amounts of antigen A and antigen B on each RBC, we observed that RBC from AB individuals differed with respect to the fluorescent staining pattern on their surface on colocalization of the A and B antigens (Figure 1).

**Figure 1:**
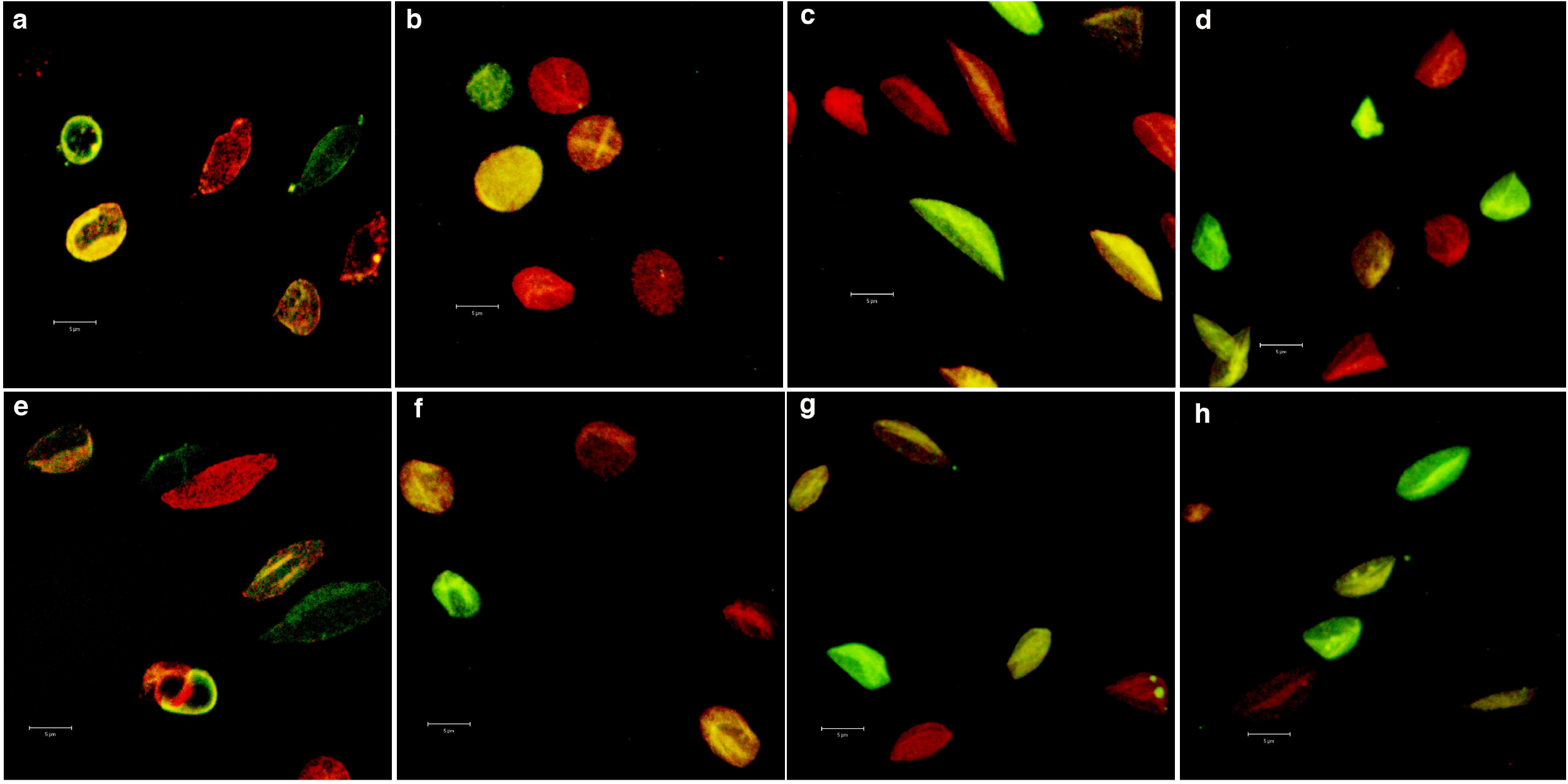
Mosaicism for antigens A and B on RBC from different subjects revealed by immunofluorescent staining: Panels **a-h** represent RBC from eight different individuals. RBC were fixed on cover slips and stained with antibodies to stain antigen A with Cy3-tag (red fluorescence) and antigen B with FITC-tag (green fluorescence). Overlay of red and green fluorescence appears as yellow fluorescence. RBC from eight subjects are shown at 100X magnification.

The samples appeared to fall into four distinct categories. In group (i), most of the RBC were stained for antigen B but differentially stained for antigen A (Figure 2, A-F). In group (ii), almost all RBC were stained for antigen A but only some cells showed staining for antigen B; in other words, staining for antigen B was differential, i.e., quantitatively different from cell to cell (Figure 2, G-L). In group (iii), RBC appeared to be stained differentially for both antigens (Figure 2, M-R). Finally, in group (iv), all the RBC stained equally for both antigens (Figure 2, S-X).

**Figure 2:**
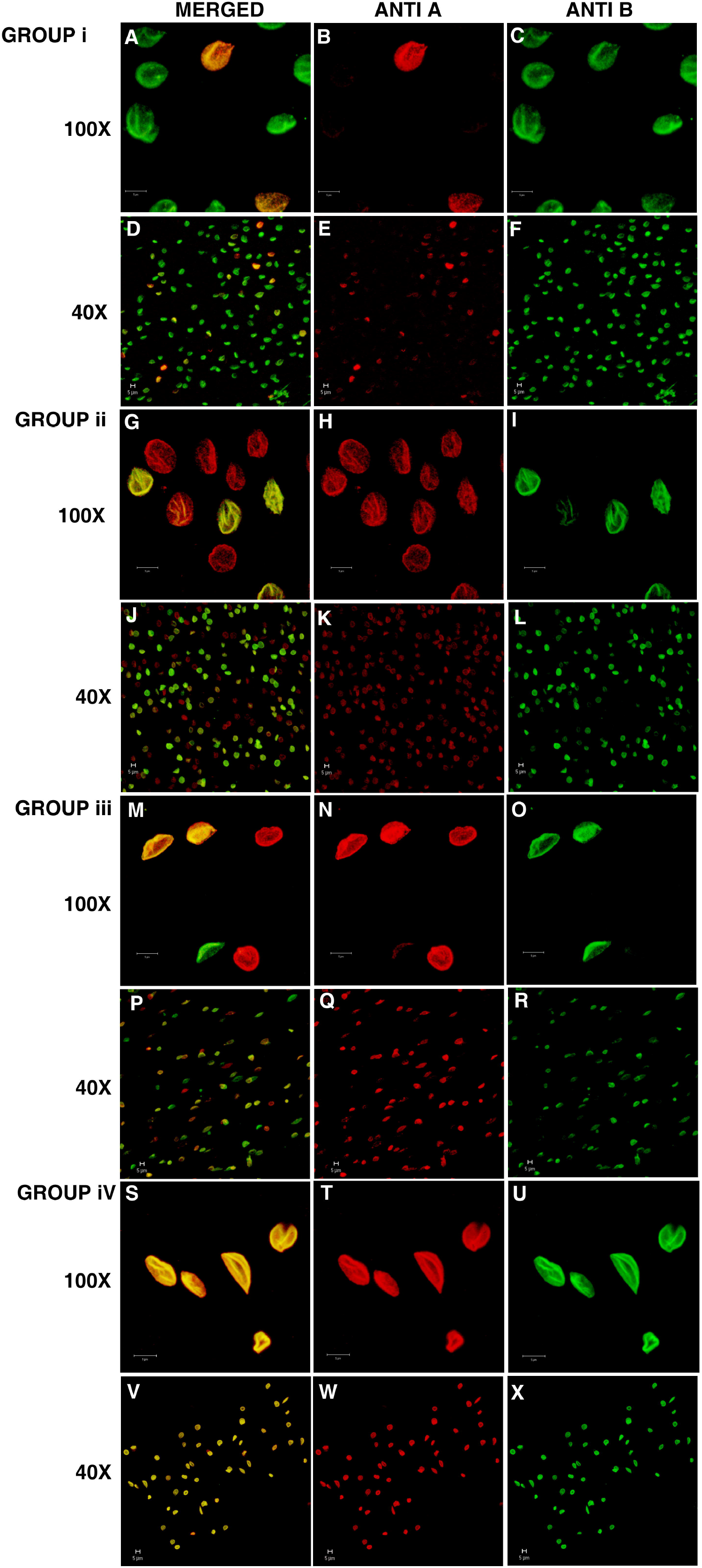
Immunofluorescent staining for antigens A and B on RBC from individuals of AB blood group: RBC were fixed on cover slips and stained with antibodies to stain antigen A with Cy3-tag (red fluorescence) and antigen B with FITC-tag (green fluorescence). Left panel shows co-localization of antigens A and B (yellow fluorescence); middle panel shows staining for antigen A; and right panel shows staining for antigen B. Cells from representative samples from each category are shown at 100X and 40 X magnifications. A-F: cells from group (i) - stained predominantly for antigen B and differential staining for antigen A; G-L: cells from group (ii) - stained predominantly for antigen A and differential staining for antigen B; M-R: cells from group (iii) - differential staining for both antigens; S-X: cells from group (iv) - equal and uniform staining for both antigens.

### Distribution of A and B antigens on RBC from AB blood group individuals

To determine the extent of expression of the antigens on individual cells, we quantified the intensity of fluorescence of FITC and Cy3 from projected confocal images using the LSM-FCS software. These RBCs were studied with respect to the number of cells showing differences and the quantitative variation between the two antigens from single cells. The data was analyzed to check for statistically significant differences, if any. The fluorescence measurements of the A and B antigens were checked for normal distribution, using the Quantile-Quantile (Q-Q) plots, which plot the distribution of the observed values against an expected normal distribution. The Q-Q plots show that the fluorescence did not show a normal distribution for most of the samples (Figure 3). Of the 24 individuals only 4 displayed normal distribution for both antigens A and B on their RBCs. On the other hand, 8 individuals showed a normal distribution for A alone, 1 individual showed a normal distribution for Antigen B, whereas a majority of 11 individuals, failed to display normal distribution for either A or B antigens on their RBCs.

**Figure 3.**
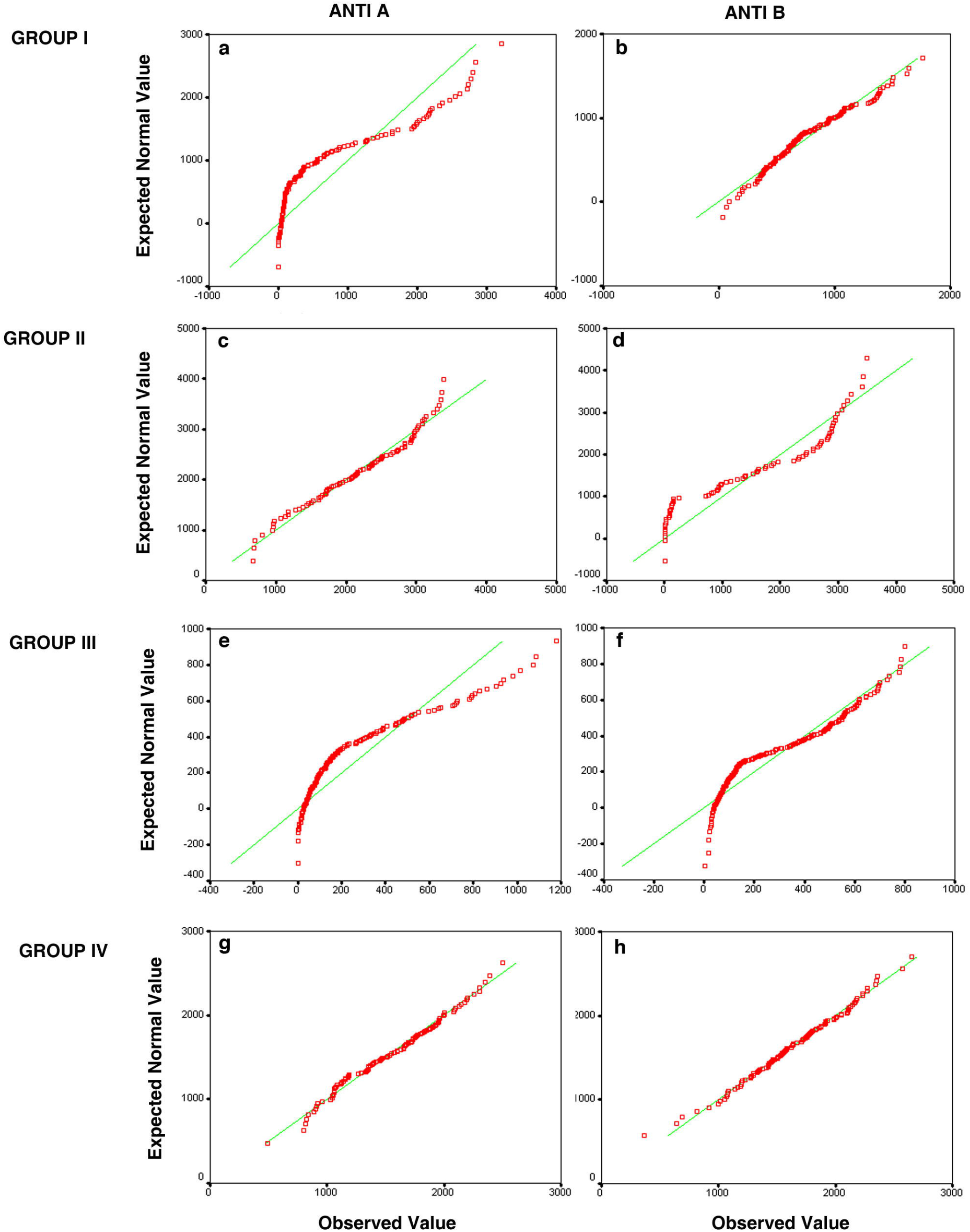
Figure shows the Q-Q plots depicting the distribution of A and B antigens from four representative individuals. The Panels **a** and **b** show the distribution of antigen A and B respectively of an AB individual who shows a differential expression of A antigen. The antigen A shows a distribution away from the normal (**a**), and antigen B (**b**) shows a normal distribution. Panels **c, d** show the distribution of an individual with differential expression of antigen B, which shows a skewed distribution **(c)** and antigen A shows a normal distribution **(d).** The panels **(e)** and **(f)** display skewed distribution of both A and B antigens in an individual who shows differential expression of both the antigens. Panels **(g)** and **(h)** show normal distribution of both the antigens A and B in an individual with equal expression of both the antigens. The Q-Q plot plots the expected normal distribution on the Y axis against the observed values on the X axis.

These data were also plotted as a box plot to show the distribution of fluorescence for antigen A (red bars) and antigen B (green bars) (Figure 4). Using the Shapiro Wilk test to assess for normality of the distribution, we found that the samples could again be assigned to 4 groups: (i), where only antigen B shows a normal distribution (subject # 19); (ii), where only antigen A shows a normal distribution (subjects #2, 3, 5, 7, 13, 17, 20 and 25); (iii), where neither antigen shows a normal distribution (subjects # 9, 11, 15, 18, 1, 24, 6, 12, 16, 23 and 26) and (iv), where both antigens show a normal distribution (subjects # 4, 8, 14, and 21).

**Figure 4:**
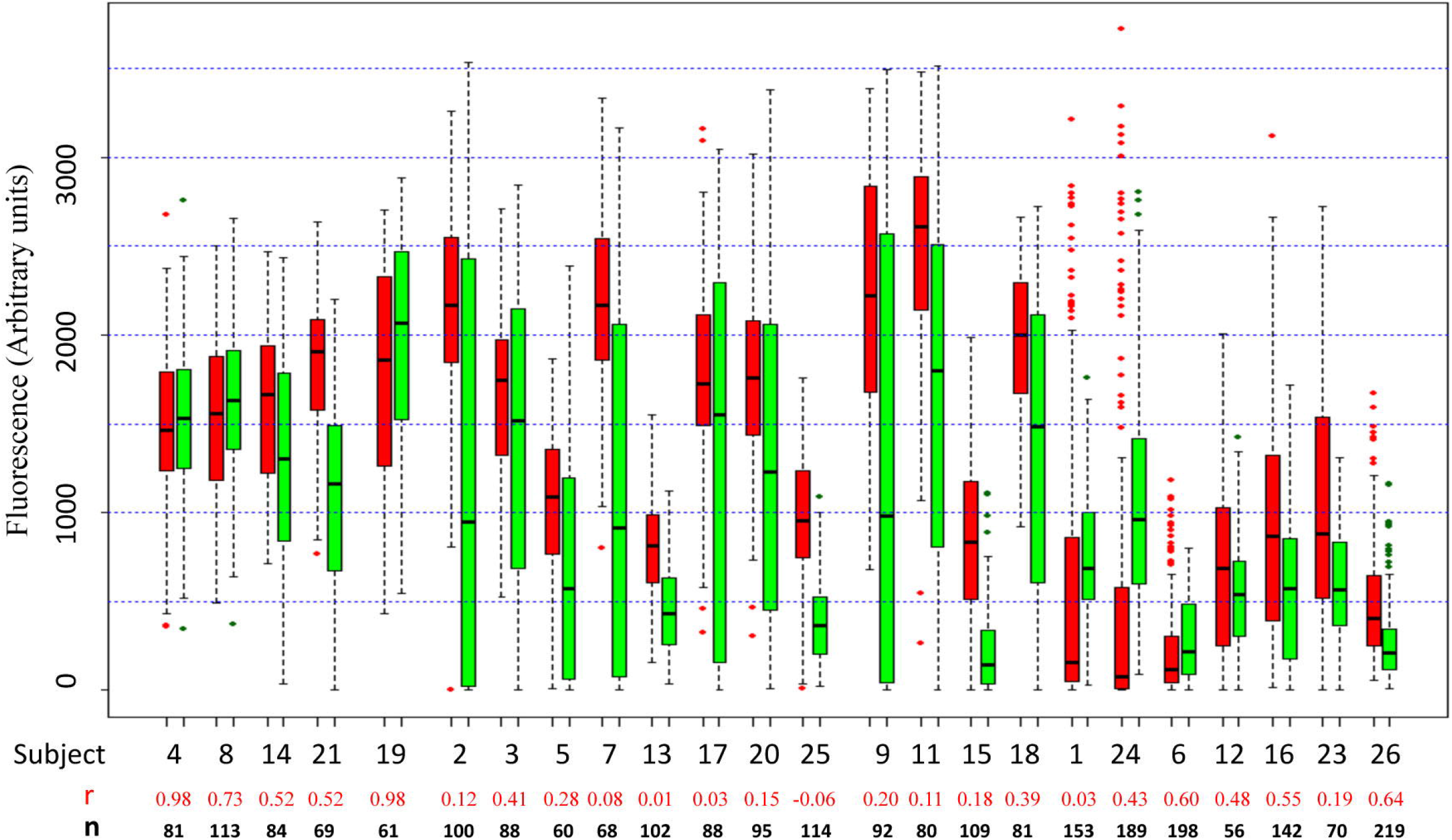
Comparison of distribution of fluorescence (arbitrary units) corresponding to antigens A and B on RBC: Box plot showing distribution for fluorescence intensity for antigens A (red boxes) and B (green boxes) stained with Cy3- and FITC-tagged antibodies. The value of Karl Pearson coefficient of correlation (r) and number of RBC quantified (n) is shown for each subject. Outliers were identified using the boxplot function of R. The box marks the limits of the upper and lower quartiles, and the median is indicated as a line within the box. Whiskers extend to 1.5*IQR (interquartile range). Filled circles indicate outliers.

Another interesting observation was that most samples showed no correlation between the amounts of antigens A and B on individual cells. As noted in Figure 4, only 2 samples (subjects # 4 and 19) showed a value above 0.9 for Karl Pearson correlation coefficient (r), and 3 other samples (subjects # 8, 6, and 26) had r>0.6.

### Range and ratio of expression of A and B antigens on RBC from AB blood group individuals

Considering that these two antigens are expressed from alleles on homologous chromosomes, this lack of correlation indicates that the expression of these antigens is not under stringent control. The extent of wide variation in the relative amount of antigens A and B on individual RBC is evident from a box plot of the ratio of fluorescence from antigen A to that from antigen B (Figure 5). If these two alleles were expressed at comparable amounts on all cells, we would expect to see a narrow range of ratios (the ratio would be 1 if the fluorescence due to the two dyes – Cy3 and FITC – had molar equivalence). Yet we see the ratios spanning 5 orders of magnitude, from 1/100 to over 1000. This is consistent with essentially independent levels of expression of these antigens and/or rates of loss during the lifetime of the RBC.

**Figure 5:**
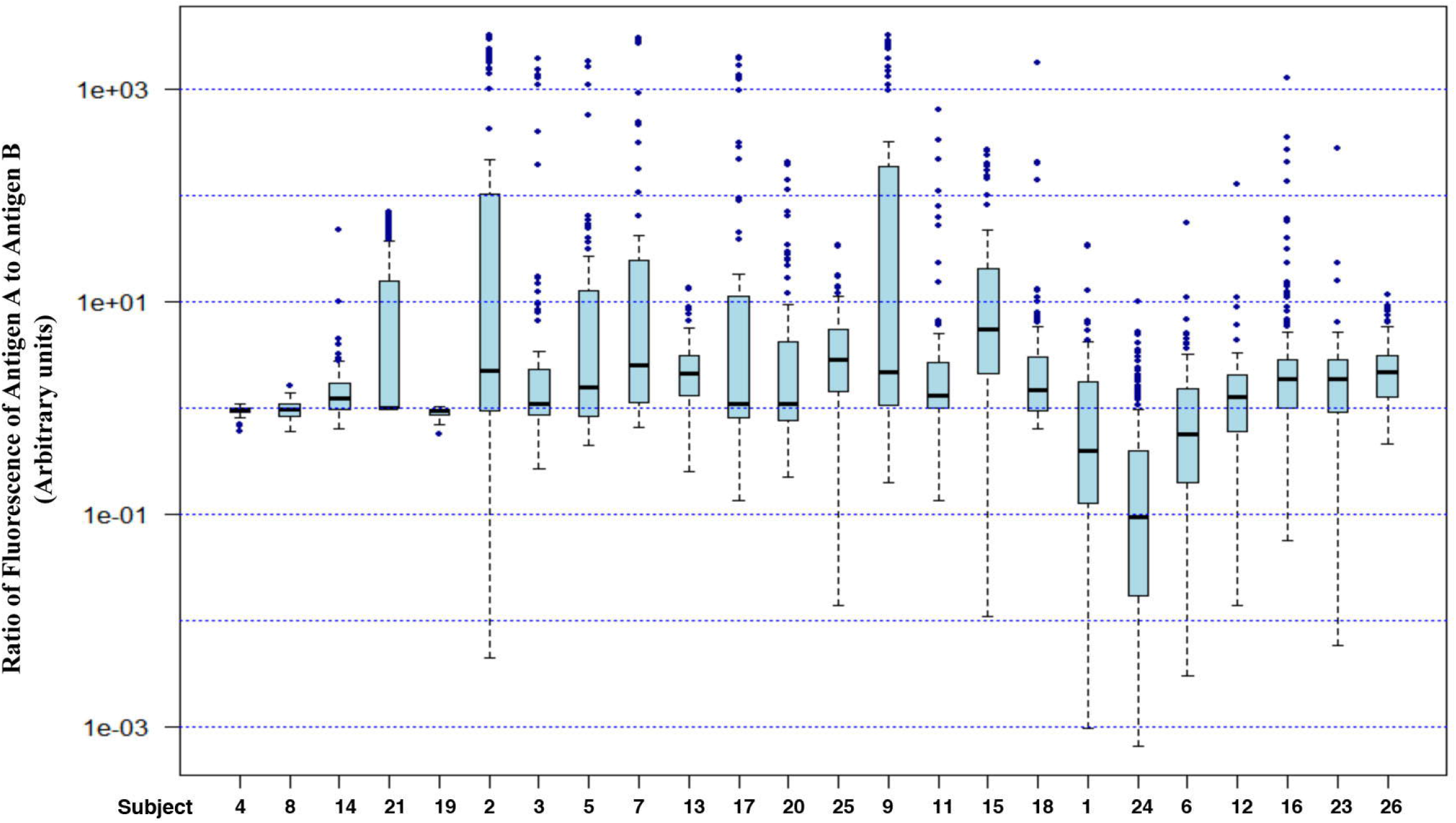
Fluorescence ratios for all individuals. Box plots and outliers were marked as for Figure 4.

Our observations are consistent with a previous report by Balea and colleagues (Balea et al., 1992) where flow cytometry of RBC from AB individuals revealed distinct sub-population of cells with equal concentrations of antigens A and B, or with maximum concentration of antigen A and minimum concentration of antigen B, or with maximum concentration of antigen B and minimum concentration of antigen A. The use of scanning fluorescence microscopy in our study has revealed more layers of heterogeneity within population of RBC in individuals of AB blood group. In many individuals, the level of heterogeneity is so high that it appears that no two RBC are alike with respect to antigens A and B on their surface. This degree of mosaicism appears to be unparalleled in human tissues.

In this cohort of 24 subjects, 12 showed normal distribution for antigen A while only five showed normal distribution for antigen B. This was an unexpected finding: differences have been reported in the promoter strength as well as the level of transcripts from various I^A^ and I^B^ alleles, but one would expect that the products of strongly or weakly expressed alleles would show a normal distribution nevertheless, even if the absolute amount of antigen A was not the same as that of antigen B. There is no synthesis of new molecules of these antigens in circulating RBC, given that these cells are devoid of ER where the galactosyl transferases encoded by I^A^ and I^B^ alleles function (Moras et al., 2017). Scant expression of the A or B antigen on circulating RBC is consistent with the corresponding enzyme being fully active only in few cells; the reasons for a possible sporadic activity, however, are not clear.

### Is the ABO locus imprinted?

To probe if the non-normal distribution of antigens from the I^A^ and/or I^B^ alleles in certain individuals was potentially due to epigenetic mechanisms dependent on the parent of origin, we attempted to correlate the alleles to paternal or maternal allele origin. The blood group of the parents of some of the subjects in this test group was known. Comparison of the blood group of the parents with the test individuals showed that there was no relationship between the alleles showing differential expression and the parent of origin (Table 1).

**Table 1:**
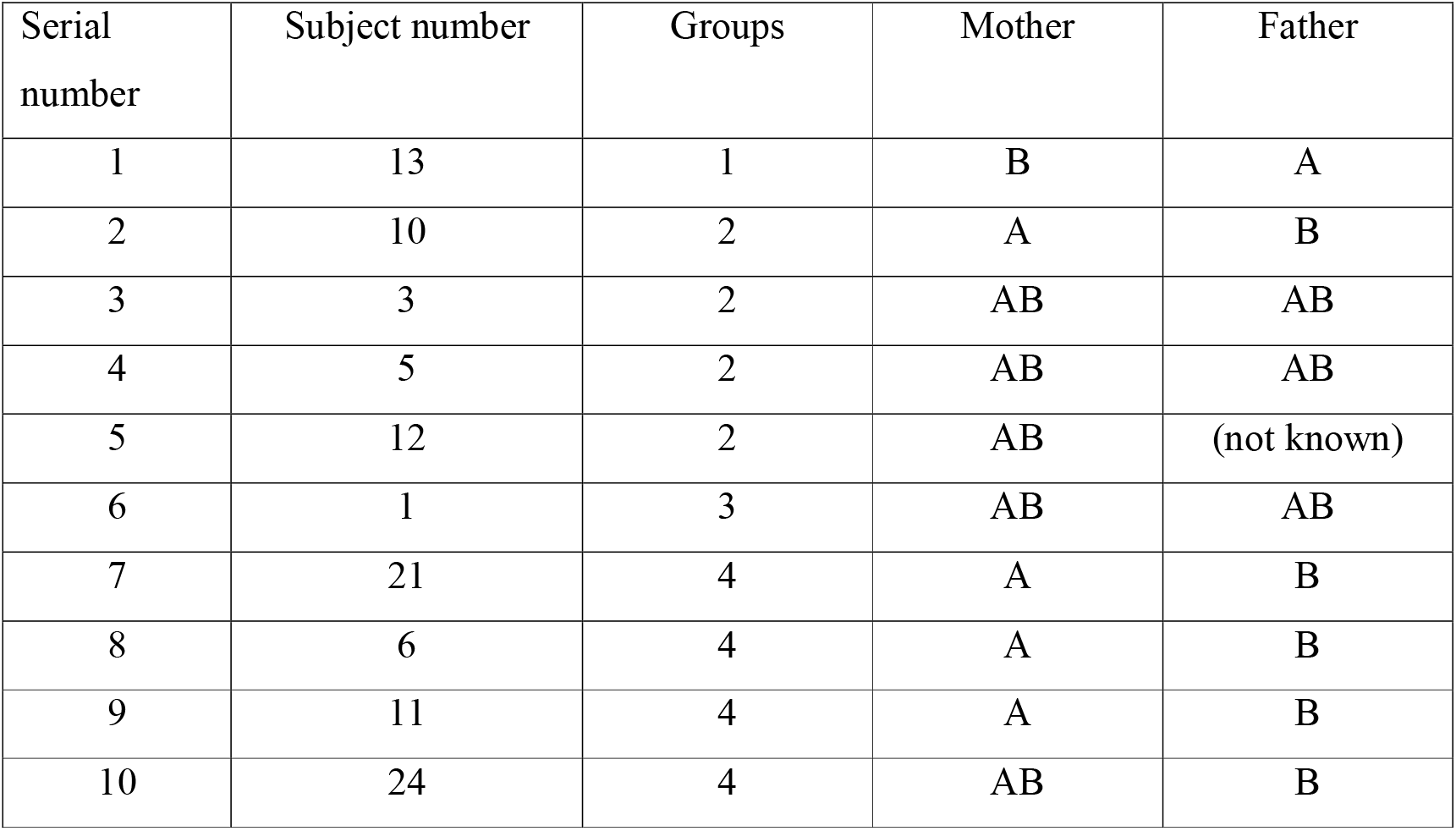
Blood groups of selected subjects and their parents. **Group 1**: Differential expression for A and Normal distribution for B; **Group 2**: Normal distribution for A and Differential expression for B; **Group 3**: Both A and B show Differential expression; **Group 4**: Both A and B show Normal distribution.

The alleles of the ABO blood group genes localizing to chromosome 9 in humans are classic examples of co-dominant gene expression (Bennett et al., 1995). Earlier studies in the lab reported a difference in the occurrence of sister chromatid exchanges on the chromosomes of *Muntiacus muntjac* (Rachel et al., 1992). More recent studies show that indeed there are a number of genes, which show allelic variation in gene expression within an individual, elucidating the fact that this could be a more common phenomenon than envisaged (Reinius and Sandberg, 2015). Allele-specific differences in gene expression have been reported and currently this concept is more widely accepted. Epigenetic factors such as single nucleotide polymorphisms in regulatory elements, parent of origin imprinting, differential methylation etc. is believed to effect such differences in gene expression (Crowley et al., 2015). Variation in transcription has been observed within cells of a population. Yet the variability at the cytoplasmic level is tightly regulated by the nucleus(Battich et al., 2015). In the RBC of the AB blood group, we observe a variation in the protein levels. The long-term follow-up of few subjects over a span of 6-7 months shows that the expression pattern of these antigens appears to be fixed for a given individual (data not shown).

The mechanisms responsible for cell-to-cell variations in A and/or B expression is not clear. Transcription from the ABO locus is under multiple regulatory elements besides its constitutively active proximal promoter (Hata et al., 2002), which include, distal enhancer elements that display binding of cell type specific factors, such as hematopoietic-specific GATA-1/2, and RUNX1 (Sano et al., 2012), or Elf2 (Sano et al., 2016) in epithelial cells. ABO locus-associated mini-satellite repeats(Irshaid et al., 1999) and CpG islands (Kominato et al., 1999) further regulate transcription from this locus. Genome-wide analysis of allele-specific expression has revealed various mechanisms that result in allelic-mosaicism in a tissue, such as, position effect variegation due to variable heterochromatin formation, epigenetic modifications such as methylation (Nag et al., 2013), hyper-mutation, cellular volume and number of mitochondria within a cell (Eckersley-Maslin and Spector, 2014; Battich et al., 2015). Interestingly, recent studies have highlighted temporally-regulated transcriptional bursts from individual alleles (Suter et al., 2011; Larsson et al., 2019) such that, such tandem events could result in differential allele expression. However, since the matured RBCs lack DNA, it is not possible to determine if the sporadic or non-coordinated transcription of the alleles is due to any of these mechanisms or a novel pathway, initiated in erythropoetic progenitor cells. Since the A and B antigens are expressed on most cell types (Clausen and Hakomori, 1989) it is possible that characterization of clones from single cells taken from some other somatic tissues may reveal modifications in the genotype that can be correlated to the phenotype, however, it is beyond the scope of this study.

It is also not clear whether this heterogeneity has any physiological significance. A variety of functions have been proposed for the ABO antigens; certain blood types have been associated with increased risk for certain medical conditions, or infections (Reviewed in (Liumbruno and Franchini, 2013; Cooling, 2015)). The revelation of extensive heterogeneity in the quantitative profiles of these antigens suggests that the use of quantitative immunofluorescence instead of the traditional method of agglutination reactions to determine the blood group of individuals may allow more thorough correlation of these risk factors with specific antigen profiles.

The differential and combinatorial expression from autosomal homologues, can give rise to cell-specific expression, not only giving each cell a unique signature, but accounting as well for individual specific gene expression. This may be one of the mechanisms by which nature can bring about phenotypic variation within a single genotype. It would be interesting to investigate if the differential response to drugs by individuals can be understood better by studying combinatorial expression of sets of genes.

## Author Contributions Statement

Conceived and designed the experiments: RAJ. Performed the experiments: AB, BS, PM.

Analyzed the data: RAJ, AB, VJ. Wrote the paper: RAJ, VJ.

## Acknowledgements

We thank Dr. R. Sarkar for help with statistics, Ms. R. Nandini for technical help with microscopy and CSIR for financial assistance to RAJ and AB.

